# Investigating the genomic landscape of novel coronavirus (2019-nCoV) to identify non-synonymous mutations for use in diagnosis and drug design

**DOI:** 10.1101/2020.04.16.043273

**Authors:** Manish Tiwari, Divya Mishra

## Abstract

Novel coronavirus has wrecked medical and health care facilities claiming ~5% death tolls globally. All efforts to contain the pathogenesis either using inhibitory drugs or vaccines largely remained futile due to a lack of better understanding of the genomic feature of this virus. In the present study, we compared the 2019-nCoV with other coronaviruses, which indicated that bat-SARS like coronavirus could be a probable ancestor of the novel coronavirus. The protein sequence similarity of pangolin-hCoV and bat-hCoV with human coronavirus was higher as compared to their nucleotide similarity denoting the occurrence of more synonymous mutations in the genome. Phylogenetic and alignment analysis of 591 novel coronaviruses of different clades from Group I to Group V revealed several mutations and concomitant amino acid changes. Detailed investigation on nucleotide substitution unfolded 100 substitutions in the coding region of which 43 were synonymous and 57 were of non-synonymous type. The non-synonymous substitutions resulting into 57 amino acid changes were found to be distributed over different hCoV proteins with maximum on spike protein. An important diamino acid change RG to KR was observed in ORF9 protein. Additionally, several interesting features of the novel coronavirus genome have been highlighted in respect to various other human infecting viruses which may explain extreme pathogenicity, infectivity and simultaneously the reason behind failure of the antiviral therapies.

**Summary:** This study presents a comprehensive phylogenetic analysis of SARS-CoV2 isolates to understand discrete mutations that are occurring between patient samples. This analysis will provide an explanation for varying treatment efficacies of different inhibitory drugs and a future direction towards a combinatorial treatment therapies based on the kind of mutation in the viral genome.

## Introduction

Current and last two decades saw emergence of zoonotic coronavirus (CoV) crossing the species barrier ultimately infecting human species resulting in pandemics such as severe acute respiratory syndrome (SARS) and Middle East respiratory syndrome (MERS) (Drosten et al., 2003; Zaki et al., 2012). An apocalyptic threat is posed by a sempiternal pathogen ruining the health and the economies on global scale. A severe pneumonia outbreak starting in December, 2019 in the Wuhan city, Hubei Province, China was caused by novel CoV referred as “2019 novel coronavirus” or “2019-nCoV” (Huang et al., 2020; Zhu et al., 2020). CoVs are RNA viruses with wide host pathogenesis in mammals including humans, pangolins and birds. Genetically the CoVs were categorised into four major genera: *Alphacoronavirus, Betacoronavirus, Gammacoronavirus*, and *Deltacoronavirus* (Li, 2016). The alpha and beta CoVs infect mammals whereas the gamma and delta CoVs infect birds (Tang et al., 2015). Primary symptoms associated with CoV infection include respiratory, hepatic, enteric and neurological diseases. Previous investigation showed that there are 6 type of CoVs (hCoV-NL63, hCoV-229E, hCoV-OC43, hCoV-HKU1, SARS-CoV, and MERS-CoV) which can infect the human species. HCoV-NL63, hCoV-229E belongs to alphaCoV genus while rest belongs to betaCoV genus. (Tang et al., 2015). The betaCoVs appears to be *prima-facie* genre of CoVs which will peril universal human civilization in upcoming decades. Recently, the 2019-nCoV outbreak spread from China to the intercontinental arena and already infected 0.3 million people globally claiming ~13000 (~4.3%) deaths till 21^st^ March 2020 (https://www.worldometers.info/coronavirus/#countries). China and Italy were the epicentres until now and chances for more calamitous centres cannot be ruled out in near future. Genome sequence analysis of SARS, MERS and 2019-nCoV confirmed its presence in betaCoVs family and divergence from the other two viruses (Zhu et al., 2020). The 2019-nCoV is a positive-strand RNA viruses with ~29 Kb genome size, 125 nm in diameter and 6 to 11 open reading frames (ORFs) (Song et al., 2019). Viral genome encodes for 4 major structural proteins namely envelope (E), spike (S), membrane (M) and 3–5 nucleocapsid (N) proteins. The genome starts with short untranslated regions (5’ UTR) followed by genes 5’-replicase (rep gene), S, E, M, N and 3’ UTR (Song et al., 2019). Two-third of the genome is represented by the rep gene at 5’ end which encodes for non-structural protein (Nsp). Spike protein is responsible for receptor binding and corresponding viral entry into the host and hence important target for future drugs to restrict the viral titre (Du et al., 2009, 2017). Viral assembly relies primarily on M and E proteins and RNA synthesis is achieved by action of N protein (Song et al., 2019).

To mitigate the severity of 2019-nCoV, researchers around the world are trying to develop antibodies and vaccine against this deadly virus. The problem with the delay in antiviral medication is superficial understanding of the virus. A dire need is to unravel the mutations in the viral genome and concomitant amino acid changes occurring presumably due to varying geographical location or upon interaction with the diverse human immune system. Various reports compared the SARS, MERS, bat and pangolin coronaviruses and paved way for significant findings, still leaving a lacunae in terms of the variations in the hCoV genomes and comparison with the previous available viruses resources. The present study deals with the mutations in the hCoV genomes and resulting change in amino acids.

## Results and Discussion

### Comparative genomic analyses of human novel coronavirus with other coronaviruses

Genomic features may provide an important clue about the relatedness and evolution of the organism. In order to get an insight into the similitude and dissimilitude between human novel coronavirus (hCoV) and other coronaviruses, the genome sequence of human novel coronavirus (hCoV) were compared with bat coronavirus (GU190215.1) (Drexler et al., 2010), severe acute respiratory syndrome-related coronavirus strain BtKY72 (KY352407.1) (Tao and Tong, 2019), bat SARS-like coronavirus isolate bat-SL-CoVZC45 (MG772933.1) (Hu et al., 2018), hCoV-19/pangolin/Guangdong/1/2019|EPI ISL 410721 (pangolin-hCoV) and hCoV-19/bat/Yunnan/RaTG13/2013|EPI ISL (bat-hCoV) which revealed approximately 81%, 81%, 89%, 90% and 96% similarity, respectively (Table 1).

**Table 1:**
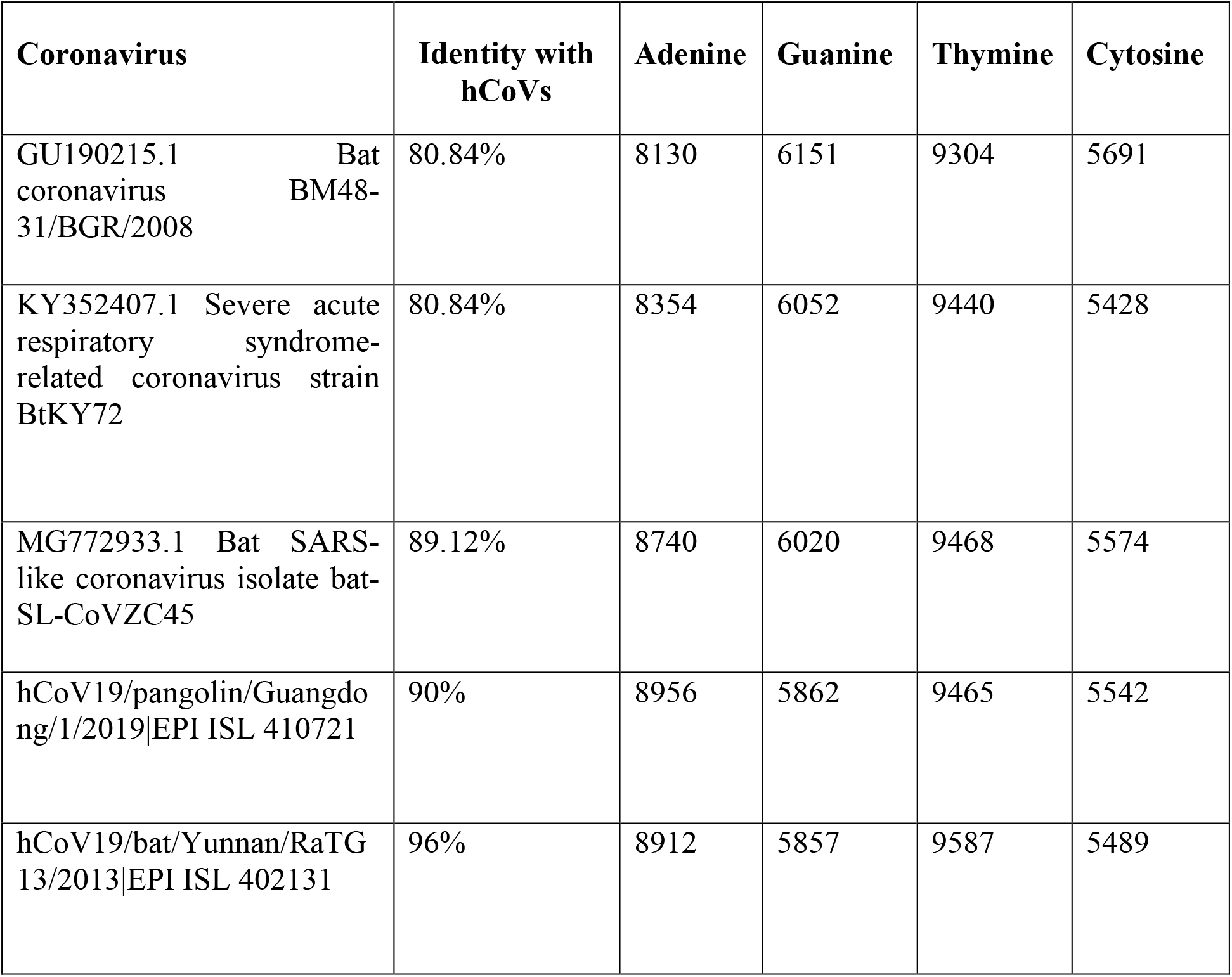
The percentage similarity of genomes of bat coronavirus, bat SARS like coronavirus, SARS coronavirus, pangolin-hCoV and bat-hCoV compared to 2019-hCoV genome

| **Coronavirus** | **Identity with hCoVs** | **Adenine** | **Guanine** | **Thymine** | **Cytosine** |
| --- | --- | --- | --- | --- | --- |
| GU190215.1 Bat coronavirus BM48-31/BGR/2008 | 80.84% | 8130 | 6151 | 9304 | 5691 |
| KY352407.1 Severe acute respiratory syndrome-related coronavirus strain BtKY72 | 80.84% | 8354 | 6052 | 9440 | 5428 |
| MG772933.1 Bat SARS-like coronavirus isolate bat-SL-CoVZC45 | 89.12% | 8740 | 6020 | 9468 | 5574 |
| hCoV19/pangolin/Guangdong/1/2019\|EPI ISL 410721 | 90% | 8956 | 5862 | 9465 | 5542 |
| hCoV19/bat/Yunnan/RaTG 13/2013\|EPI ISL 402131 | 96% | 8912 | 5857 | 9587 | 5489 |

To further assess the relationship between hCoV and other coronaviruses, alignment and phylogenetic analysis was carried out. Alignment of hCoV with above mentioned viruses showed that several nucleotide sites were unique in hCoV sequences when compared to other coronaviruses (Table S1). Among these sites, C:T (hCoV:other coronavirus) is the most prevalent substitution followed by T:C, G:A and A:G (Table S1). Many regions were absent in bat and SARS coronavirus genome when compared to hCoV, bat-hCoV and pangolin-hCoV. Among these regions, one of largest portion is of 391 nt (28026-28417) coding for ORF8 protein in hCoV putatively involved in interspecies transmission (Lau et al., 2015). Genomic similarities and alignment indicate that several mutation events over the time is responsible for emergence of human novel coronavirus. Further a phylogenetic analysis between these viruses displayed that hCoVs are closer to bat/SARS-like virus (MG772933.1) and distant from SARS coronavirus (KY352407.1) and bat coronavirus (GU190215.1) (Figure 1). These results demonstrate that SARS coronavirus and bat coronavirus (GU190215.1) could be apparent ancestor of other coronaviruses studied in the investigation.

**Figure 1.**
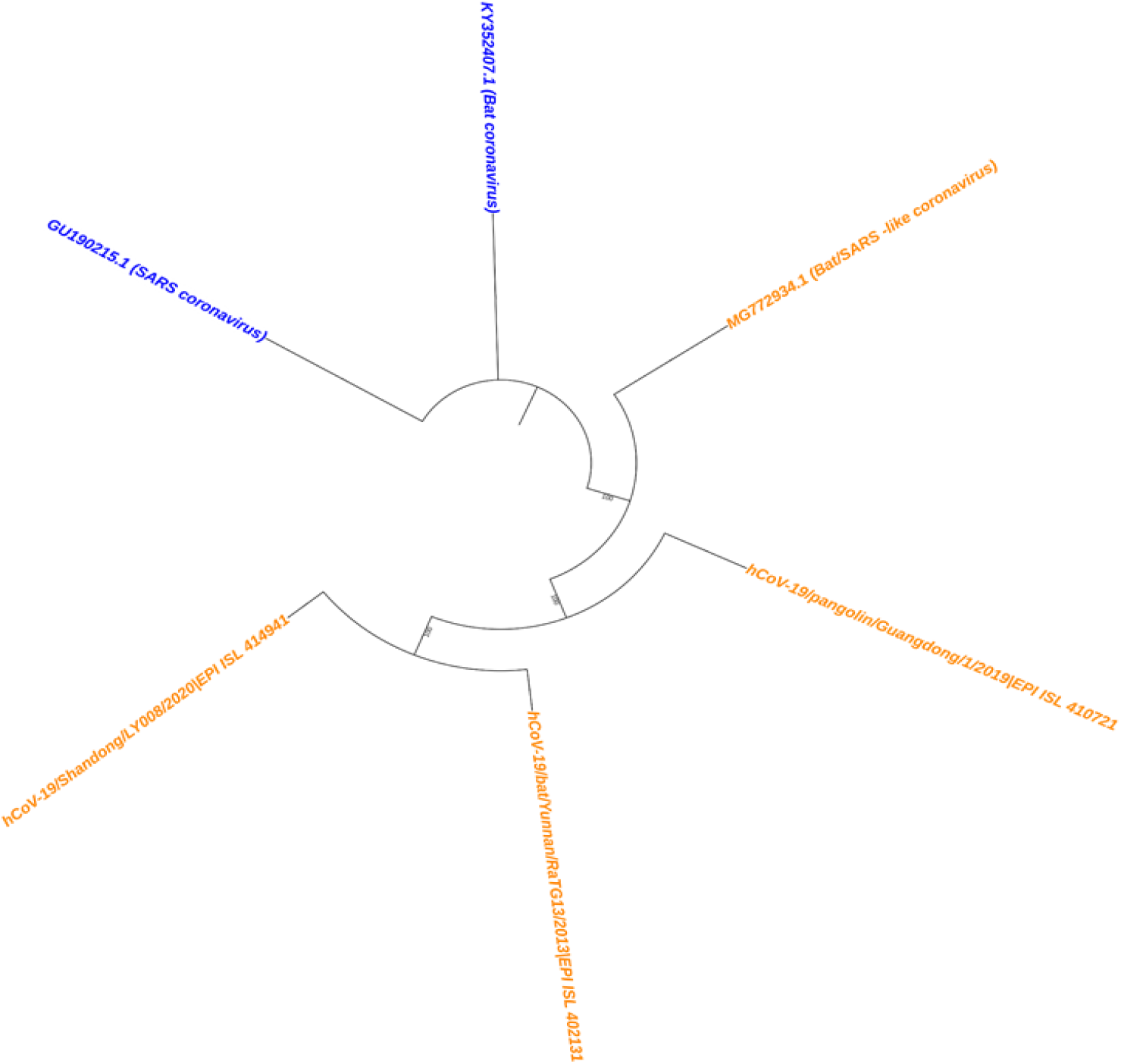
Phylogenetic relationship of hCoVs with other coronavirus. Phylogenetic analysis of bat coronavirus (GU190215.1), severe acute respiratory syndrome-related coronavirus strain BtKY72 (KY352407.1), bat SARS-like coronavirus isolate, bat-SL-CoVZC45 (MG772933.1), pangolin-hCoV, bat-hCoV and hCoV using the maximumlikelihood method (RAxML) keeping the bootstrap value 1000. Human coronavirus (hCoV, pangolin-hCoV, bat-hCoV) and bat SARS-like coronavirus falls in one clade while Severe acute respiratory syndrome-related coronavirus strain BtKY72 (KY352407.1) and bat coronavirus (GU190215.1) in another clade.

Scrutiny of nucleotide and amino acid in coding region of the genome revealed that the hCoV genome share 92.67% and 96.92% similarity at nucleotide level with pangolins and bat hCoV genome, whereas the similarity level increased up to 97.82% and 98.67% at amino acid level (Table 2). This indicates that most substitutions taking place were of synonymous type. Among various protein coding genes Nsp4-10, Nsp12-14, Nsp16, M, E and ORF6 shared highly conserve amino acid composition between bat-hCoV and hCoV with >99% similarity, especially Nsp7-10, Nsp16, E and ORF6 share 100% amino acid similarity (Table 2). The 100% similarity in these regions across 591 hCoVs, bat and pangolin-hCoV mark them to be a probable target region for future antibodies and vaccine therapy. Notably, Nsp2 and Nsp14 region in hCoVs were most diversified in terms of nucleotide when compared to pangolin and bat-hCoV, whereas ORF10 and E regions were the least diverse (Table 2).

**Table 2:**
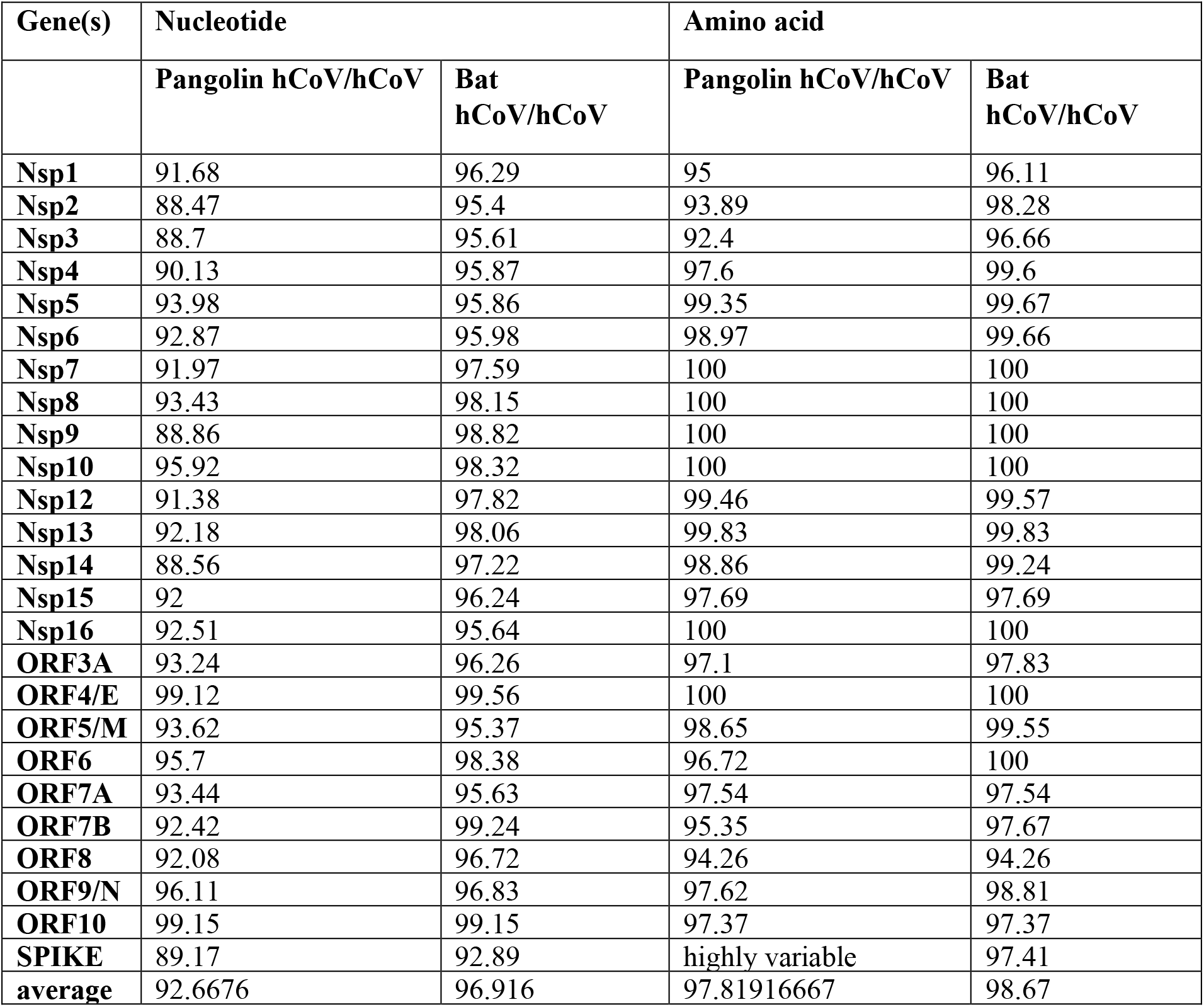
The comparison of nucleotide and amino acid similarity of pangolin-hCoV and bat-hCoV with hCoV

| **Gene(s)** | **Nucleotide** |  | **Amino acid** |  |
| --- | --- | --- | --- | --- |
|  | **Pangolin hCoV/hCoV** | **Bat hCoV/hCoV** | **Pangolin hCoV/hCoV** | **Bat hCoV/hCoV** |
| **Nsp1** | 91.68 | 96.29 | 95 | 96.11 |
| **Nsp2** | 88.47 | 95.4 | 93.89 | 98.28 |
| **Nsp3** | 88.7 | 95.61 | 92.4 | 96.66 |
| **Nsp4** | 90.13 | 95.87 | 97.6 | 99.6 |
| **Nsp5** | 93.98 | 95.86 | 99.35 | 99.67 |
| **Nsp6** | 92.87 | 95.98 | 98.97 | 99.66 |
| **Nsp7** | 91.97 | 97.59 | 100 | 100 |
| **Nsp8** | 93.43 | 98.15 | 100 | 100 |
| **Nsp9** | 88.86 | 98.82 | 100 | 100 |
| **Nsp10** | 95.92 | 98.32 | 100 | 100 |
| **Nsp12** | 91.38 | 97.82 | 99.46 | 99.57 |
| **Nsp13** | 92.18 | 98.06 | 99.83 | 99.83 |
| **Nsp14** | 88.56 | 97.22 | 98.86 | 99.24 |
| **Nsp15** | 92 | 96.24 | 97.69 | 97.69 |
| **Nsp16** | 92.51 | 95.64 | 100 | 100 |
| **ORF3A** | 93.24 | 96.26 | 97.1 | 97.83 |
| **ORF4/E** | 99.12 | 99.56 | 100 | 100 |
| **ORF5/M** | 93.62 | 95.37 | 98.65 | 99.55 |
| **ORF6** | 95.7 | 98.38 | 96.72 | 100 |
| **ORF7A** | 93.44 | 95.63 | 97.54 | 97.54 |
| **ORF7B** | 92.42 | 99.24 | 95.35 | 97.67 |
| **ORF8** | 92.08 | 96.72 | 94.26 | 94.26 |
| **ORF9/N** | 96.11 | 96.83 | 97.62 | 98.81 |
| **ORF10** | 99.15 | 99.15 | 97.37 | 97.37 |
| **SPIKE** | 89.17 | 92.89 | highly variable | 97.41 |
| **average** | 92.6676 | 96.916 | 97.81916667 | 98.67 |

### Phylogenetic analyses of human novel coronavirus

We investigated the phylogenetic analysis of 591 genomic sequences of hCoV obtained from GISAID database using RAxML methods. The phylogram was majorly divided into 5 groups based on their clade division. Bat and pangolin-hCoV were categorized in the group I and all other hCoVs were categorized in group II to V (Figure 2). Group II comprises of the human 2019-nCoV mainly belonging to different province of China. However, few exceptions were also from South Korea, Japan, Vietnam, Chile, USA, India, Belgium, Spain, Germany and France hCoVs. Group III mainly comprised of the hCoVs belonging to USA while Group IV represented the hCoVs of mixed type population belonging to several countries distributed over continents. Group V possess the hCoV from European countries along with few hCoVs of America and one from Taiwan. To understand mystery underlying the clustering pattern of the hCoVs, bat and pangolin-hCoV were used as a reference sequence to observe the nucleotide substitution in hCoV members in different groups. Interestingly, hCoV members (Group II and Group III) falling in proximity to Group I have less substitution in the genome sequences (Table S2). The T:C (GroupV-hCoV:bat-hCoV and GroupV-hCoV:pangolin-hCoV) substitution were frequent in Group V as compared to hCoV representing other groups (Table S2). The genomic signature of USA-hCoVs present in Group V is very different from USA-hCoVs of Group III. This could be indicative of differences between direct and community transmission of the virus. Member belonging to each subgroup has distinct genomic features in terms of nucleotide substitution (Table S2).

**Figure 2.**
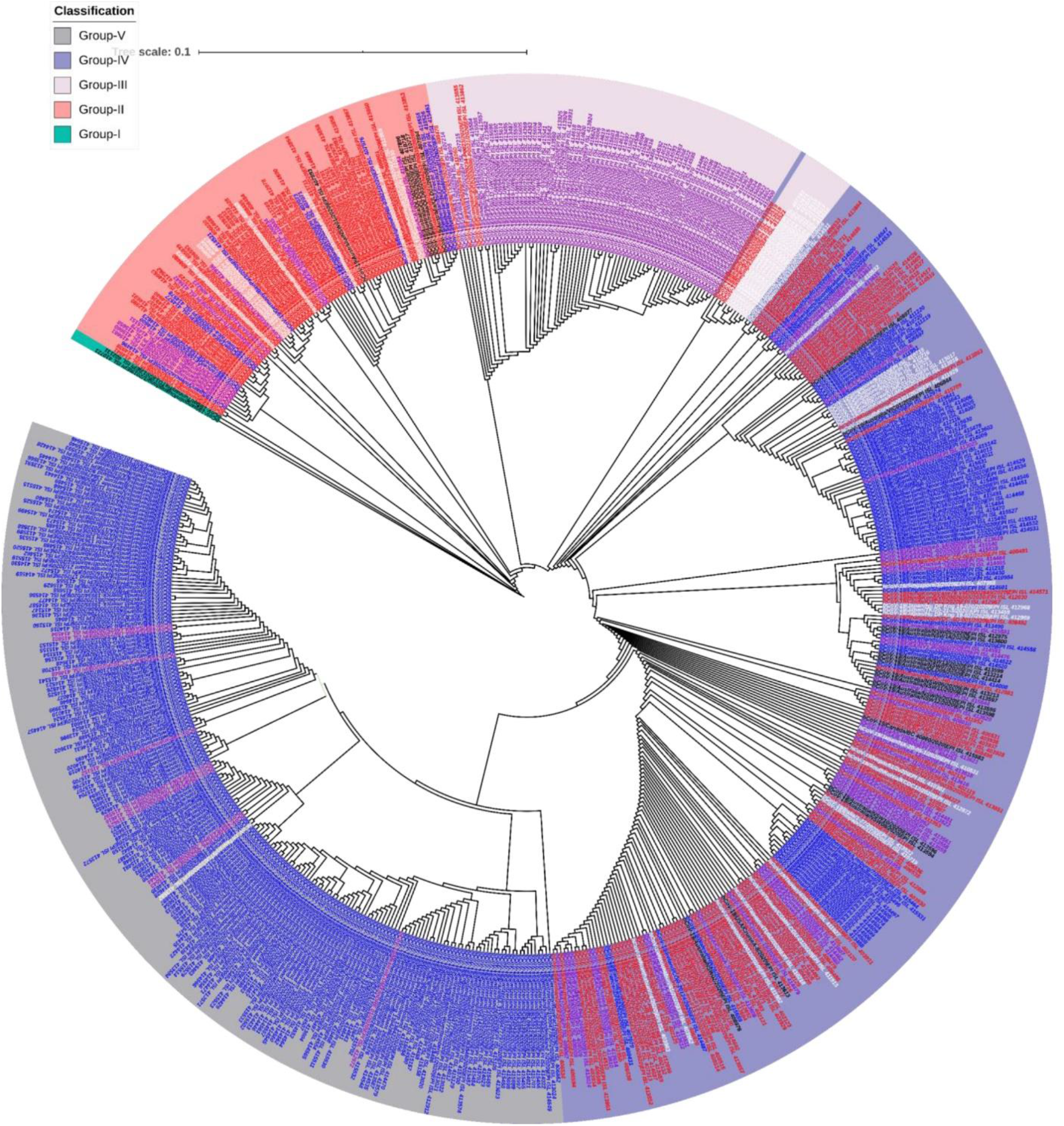
Phylogenetic relationship among 2019-nCoV. The phylogenetic tree of 519 hCoVs sequence were divided in 5 groups. Bat and pangolin hCoVs were categorised in group I and rest of the hCoVs in group II-V. The phylogram was constructed by maximum likelihood keeping the bootstrap value 1000.

### Non-synonymous substitutions and associated amino acid changes

Genomic comparison of 591 hCoV sequences among the human as well as with pangolin-hCoV and bat-hCoV revealed several sites possessing substitutions which clearly indicated the mutation in viral genome either according to the geographical locations or upon interaction with the human immune system. The nucleotide substitution in hCoV genomes were predominantly of transition type with ~45% being C:T (Figure 3A). A detailed investigation of the nucleotide substitution in the coding region of hCoVs genome with perspective of encoded amino acids revealed 43 synonymous and 57 non-synonymous substitutions (Table S3). The proteins Nsp1, Nsp5, Nsp7-10, Nsp14-16, ORF4, ORF7a, ORF7b and ORF10 mainly possessed synonymous substitutions and hence were mostly devoid of amino acid changes (Figure 3B).

**Figure 3.**
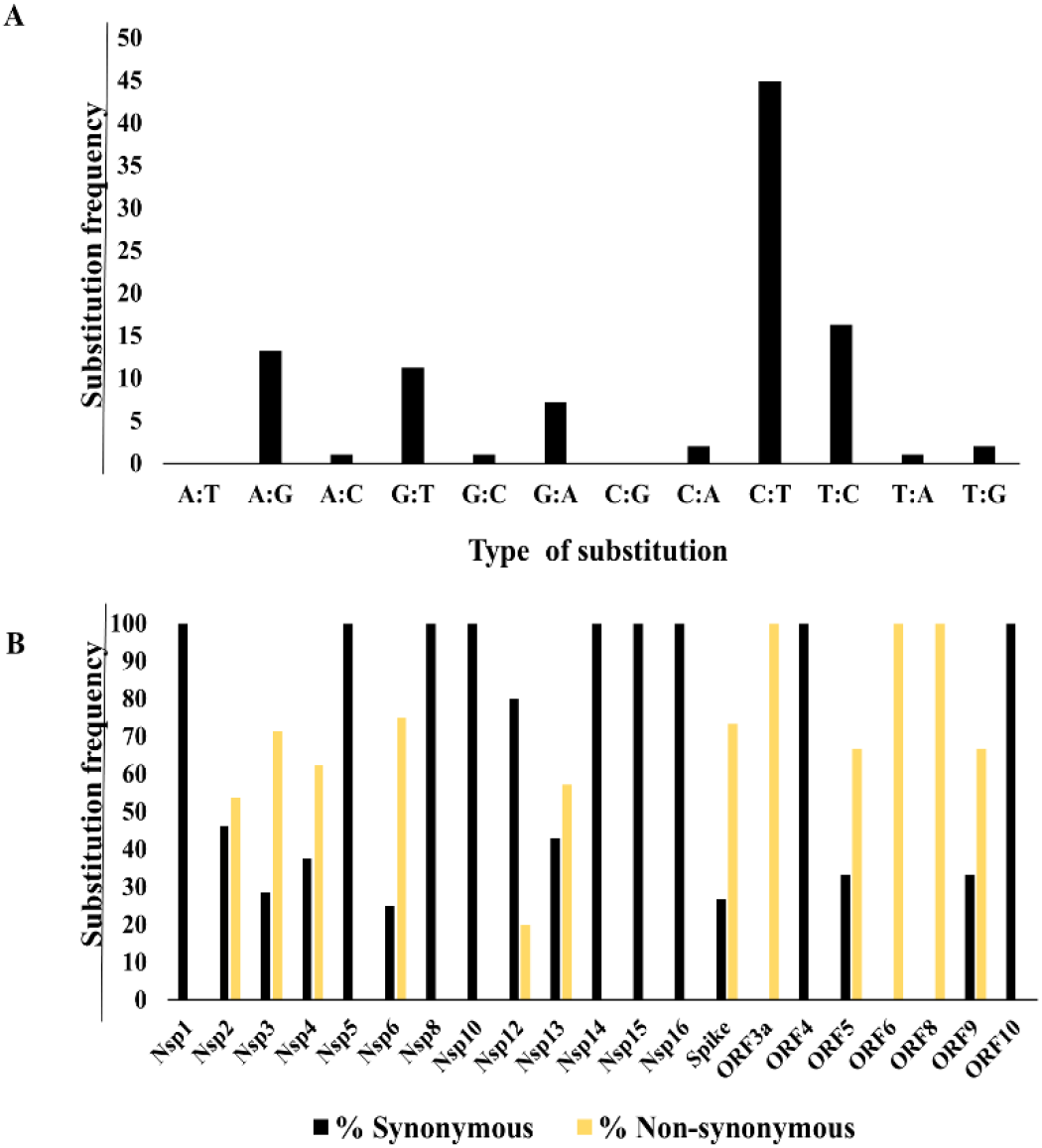
Types of substitution. **(A)** The histogram representing types of substitution. The y-axis denotes the substitution frequency and x-axis denotes the type of substitution. **(B)** The histogram representing the synonymous and non-synonymous types in various genes. The y-axis denotes the substitution frequency and x-axis denotes the region of mutation of hCoV sequence.

The 57 amino acid changes were distributed over 12 regions in the ~30kb genome. The number of amino acids substitutions varied between different regions such as 7 in Nsp2, 10 in Nsp3, 5 in Nsp4, 3 in Nsp6, 1 in Nsp12, 4 in Nsp13, 11 in Spike, 3 in ORF3a, 2 in ORF5, 1 in ORF6, 2 in ORF8 and 8 in ORF9 (Figure 4). Intriguingly, various important non-synonymous mutations were observed majorly in European and US continent while the mutations were mostly synonymous in Asian continent. These interesting observations can be used to infer the reason behind larger infectivity and pathogenicity in these regions (Table S3). Further two type of amino acid change *viz*., conservative and radial replacements were intensively studied with respect to previous reports stating the effect of such changes on the enzymatic activities. Mutations were most prevalent in the spike region followed by Nsp2, Nsp3 and ORF9 (N) (Table S3). Spike region determines the specific binding to host receptor and initiation of viral replication. This region is reported to be the most potent and indispensable for viral attachment and entry into host system. The RRAR amino acids found only in the human CoVs spike region has proved to be essential for binding to host receptor (Walls et al., 2020). We observed similar region in the hCoV genomes studied (23713-23724 region in nucleotide alignment), although there was mutation in two hCoV-England nucleotide sequences (CTCC**G**CGGCGGG in place of CTCC**T**CGGCGGG) but the resulting amino acid remained same in all hCoV genomes. These findings corroborate the essentiality of RRAR sequence for viral infection to host system. We found different type of mutation in hCoV spike protein at different places such as leucine to valine (L8V), glutamine to histidine (Q675H and also found in ORF3a:Q57H), glutamine to lysine (Q239K) and aspartate to glycine (D614G and also found in ORF5, D3G) might have potential role to augment viral infection (Table S3). Previous investigations showed mutations such as leucine to valine change in retroviral envelope protein, glutamine to lysine in influenza virus, glutamine to histidine and aspartate to glycine in H1N1 had a severe impact in virus entry, replication and cross infectivity to other species (Côté et al., 2012; Glinsky, 2010; Yamada et al., 2010).

**Figure 4.**
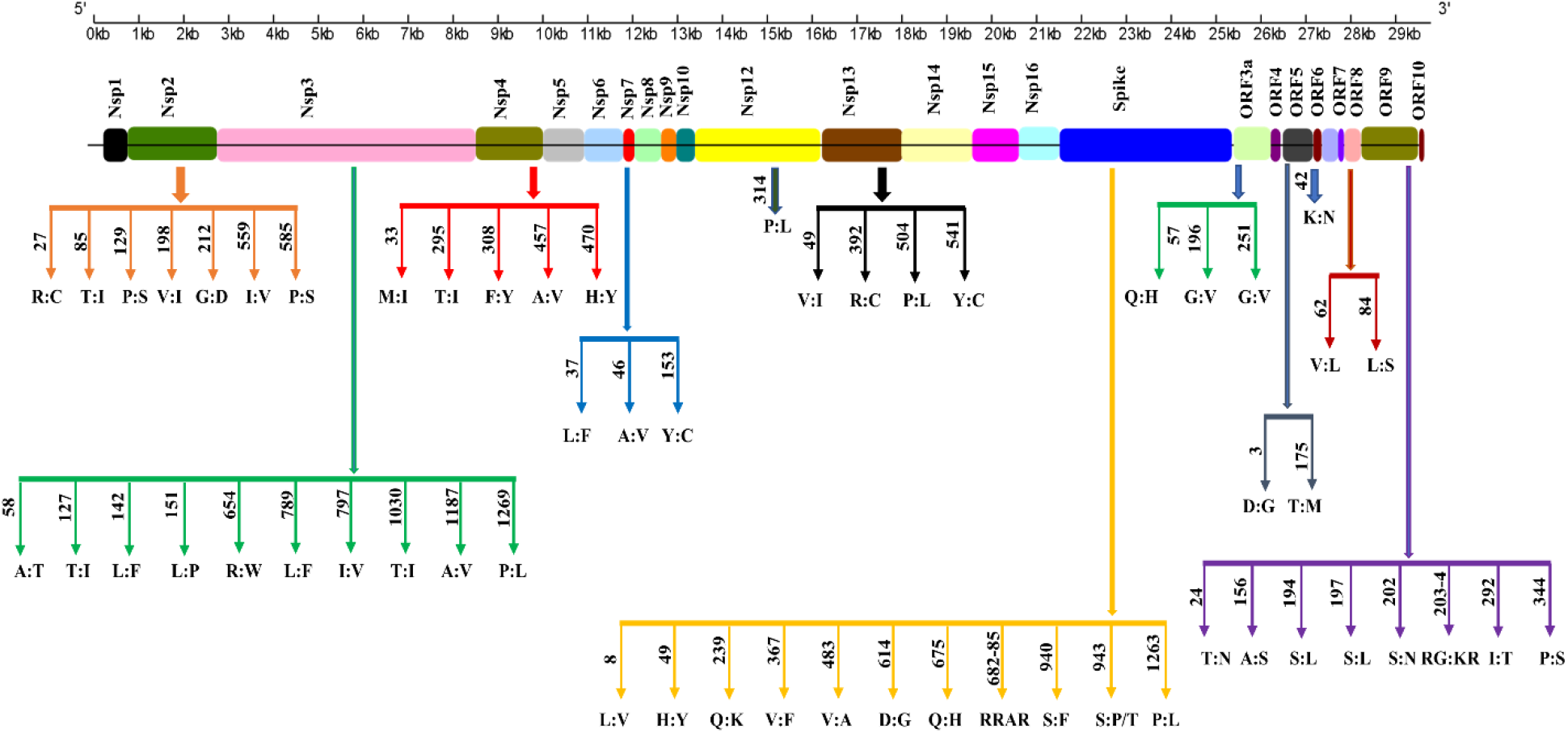
The genomic landscape of the hCoV genome representing amino acid changes. The non-synonymous mutations resulting into 57 amino acid changes in respective proteins are marked. The numerals on the arrow represent the position corresponding to amino acid change.

Additionally, mutations were present in structural proteins such as, glycine to valine mutation in ORF3a (G196V and G251V). Similar amino acid change imparts resistance against inhibitor drug saquinavir in the human immunodeficiency virus type 1 (HIV-1). This might provide an explanation why drugs used for treating HIV became a failure in case of hCoV infection (Hong et al., 1997). Notably, in ORF9 region the nucleotide sequence GGG changed to AAC in European and American continent resulting in a change of amino acid from RG to KR (AGGGGA coding for RG changed to AAACGA coding for KR, 28993-28995 in nucleotide alignment).

Furthermore, several amino acid changes were also observed in the non-structural proteins (Nsp) of the hCoVs which may affect the virulence and titer. Threonine to isoleucine substitution was observed in different Nsp proteins (Nsp2:T85I, Nsp3:T127I and T1030I and Nsp4:T295I) mainly in European and US samples. Earlier reports established that threonine to isoleucine substitution increased viral infectivity of Ebola virus and resistance to ganciclovir in human cytomegalovirus (Kurosaki et al., 2018; Wolf et al., 1995). Importantly, alanine to valine substitution in non-structural protein, NS2A in Zika virus affects viral RNA synthesis and results in vivo viral attenuation (Marquez-Jurado et al., 2018). This mutated virus also induce a comprehensive protection against lethal challenge proposed by the wild type Zika virus. Falling in similar lines alanine to valine substitutions in non-structural proteins (Nsp3, A1187V, Nsp4, A457V and Nsp6, A46V) could reduce viral lethality of hCoVs (Table S3). These mutations might pave way towards identification of less lethal strains and help to raise immunity to counteract the noxious strains. An isoleucine to valine mutation (Nsp2, I559V and Nsp3, I797V) and methionine to isoleucine (Nsp4, M33I) were observed in hCoVs. Change of isoleucine to valine in polymerase subunit PB2 of influenza virus resulted in critically enhanced activity of reconstituted polymerase complex (Rolling et al., 2009) and M to I substitution in HIV-1 reverse transcriptase imparted resistance to nucleoside analog 2’,3’-dideoxy-3’thiacytidine (3TC) (Julias et al., 2004). Interestingly, presence of a non-synonymous substitution in RNA Dependent RNA Polymerase (RDRP) region in majority of European hCoV samples resulted in change of amino acid from proline to leucine (P314L). It will be quite interesting to validate the effect of this substitution on RDRP activity as one of the previous study established that similar change of proline-to-leucine substitution (P236L) of HIV-1 reverse transcriptase, imparts resistance against a highly specific inhibitor bisheteroarylpiperazines (BHAPs) (Fan et al., 1995). These examples clearly show that amino acid changes may significantly affect the functional competency of polymerase and the associated subunits.

In conclusion, present study enlightens about several types of mutation such as deletion, insertion and substitutions present in 2019-nCoV samples. These mutations may vary at different geographical distribution or interaction with different host systems. Few mutations also resulted in change of amino acid which may provide an explanation for failure of previously employed antiviral therapies. This research will better equip the researchers to utilize the mutated amino acid information for drug targets in particular geography and less cases of failure. Beside the substitution resulting into transformation to a more virulent strain there are number of highly conserved regions in the hCoV genome which can be used as target for inhibitory drugs and vaccine development for a large repertoire of strains. Finally, we believe that our data provide useful information pertaining the changes in genomic and proteomic features which could serve as a guide to design the future antiviral therapies and diagnostics.

### Star Methods

To analyse the phylogenetic relationship between different coronaviruses, 591 genomes were downloaded from Global Initiative on Sharing All Influenza Database (GISAID) (https://www.gisaid.org/). The hCoV is an RNA virus and the deposited sequences are in DNA format. To prevent anomaly in the data represented, complete genomes and only high coverage datasets were utilized. The genomic sequences were aligned using MUSCLE program (v3.8.31) (Edgar, 2004). The alignments were utilized to deduce various nucleotide substitutions and maximum likelihood phylogenetic tree with 1000 bootstrap was constructed by RAxML program (Stamatakis, 2014). The alignment and tree were visualized using Jalview 2.11.0 (Waterhouse et al., 2009) and iTOL respectively (Letunic and Peer, 2007). Different substitutions and resulting amino acid changes were analyzed between human, bat, pangolin and SARS coronavirus genomes. To deduce a mutation or amino acid change only those confirmed in three individual genomes were considered (replicates for biological significance).

## Acknowledgements

We kindly acknowledge National Institute of Plant Genome Research (NIPGR) and Department of Biotechnology, Govt. of India (http://www.dbtindia.nic.in).

## Author contributions

M.T. performed the computational analysis, D.M. prepared all the figures and tables. M.T. and D.M. designed the project and wrote the article.

---

**Table S1:**
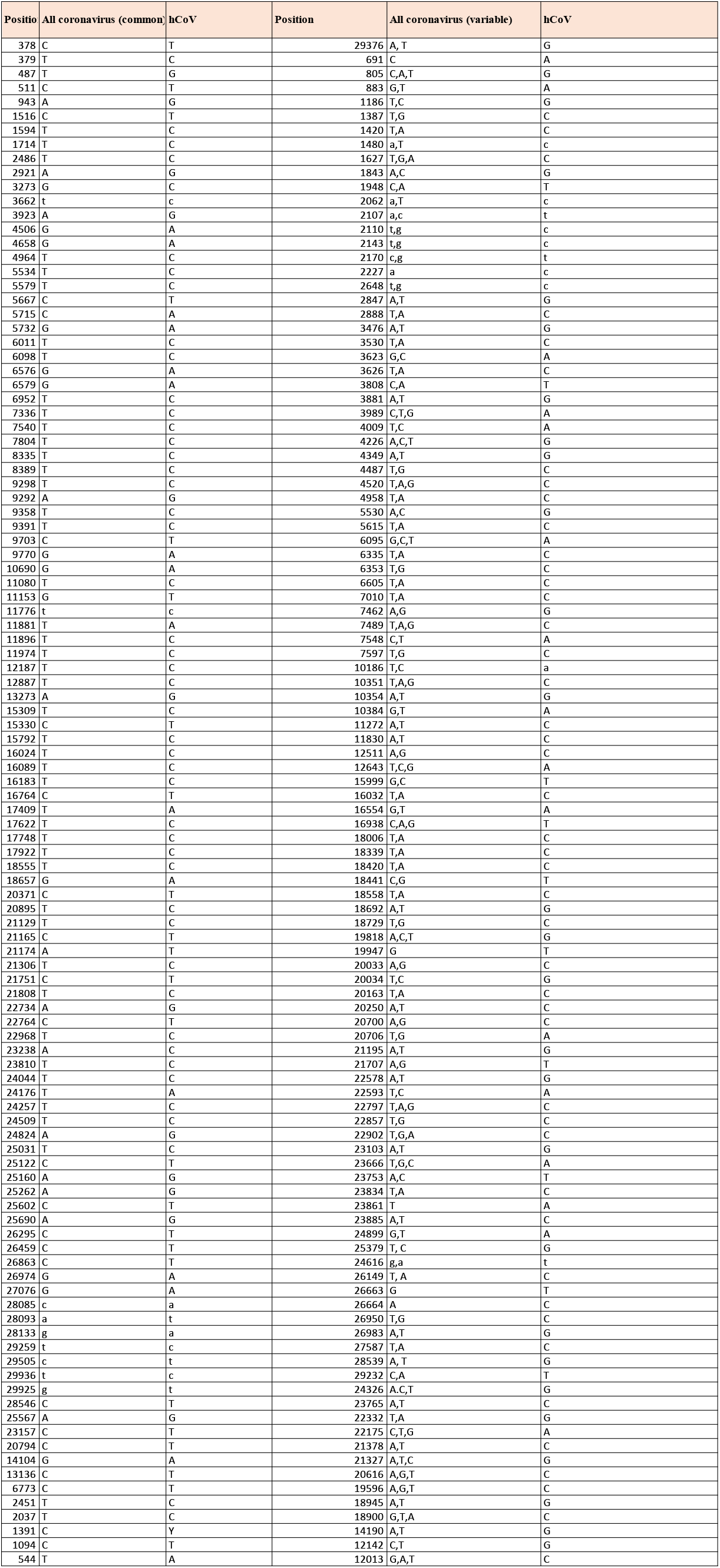
The list of subsitution in hCoV sample (CoV-19/Shandong/LY008/2020|EPI_ISL_414941/1-29868) with respect to all coronavirus [(found common nucleotide and variable nucleotide in SARS (KY352407.1 severe acute respiratory syndrome-related coronavirus strain BtKY7O), batCoV (GU190215.1 Bat coronavirus BM48-31/BGR/2008), bat/SARS-CoV MG772933.1 (bat SARS-like coronavirus isolate bat-SL-CoVZC45), pangolin CoV (hCoV-19 pangolin Guangdong/1/2019 EPI ISL 410721) and bat CoV (hCoV-19/bat/Yunnan/RaTG13/2013|EPI ISL 402131]

**Table S2:**
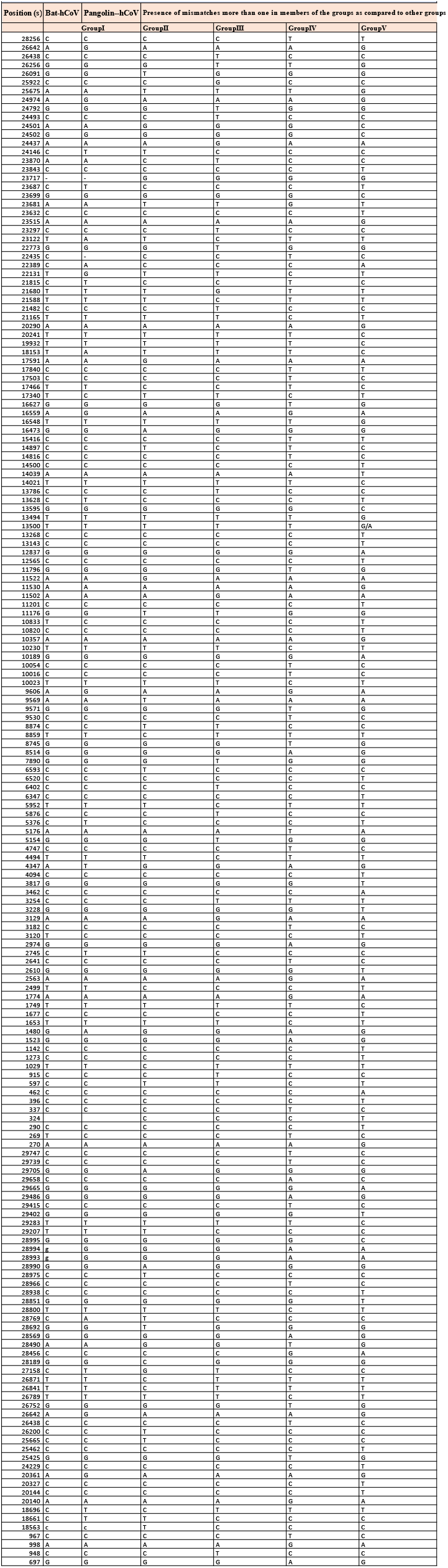
The list of nucleotide substitution in Group II-V compared to Bat-hCoV and Group II-V compared to Pangol in-hCoV.

**Table S3:**
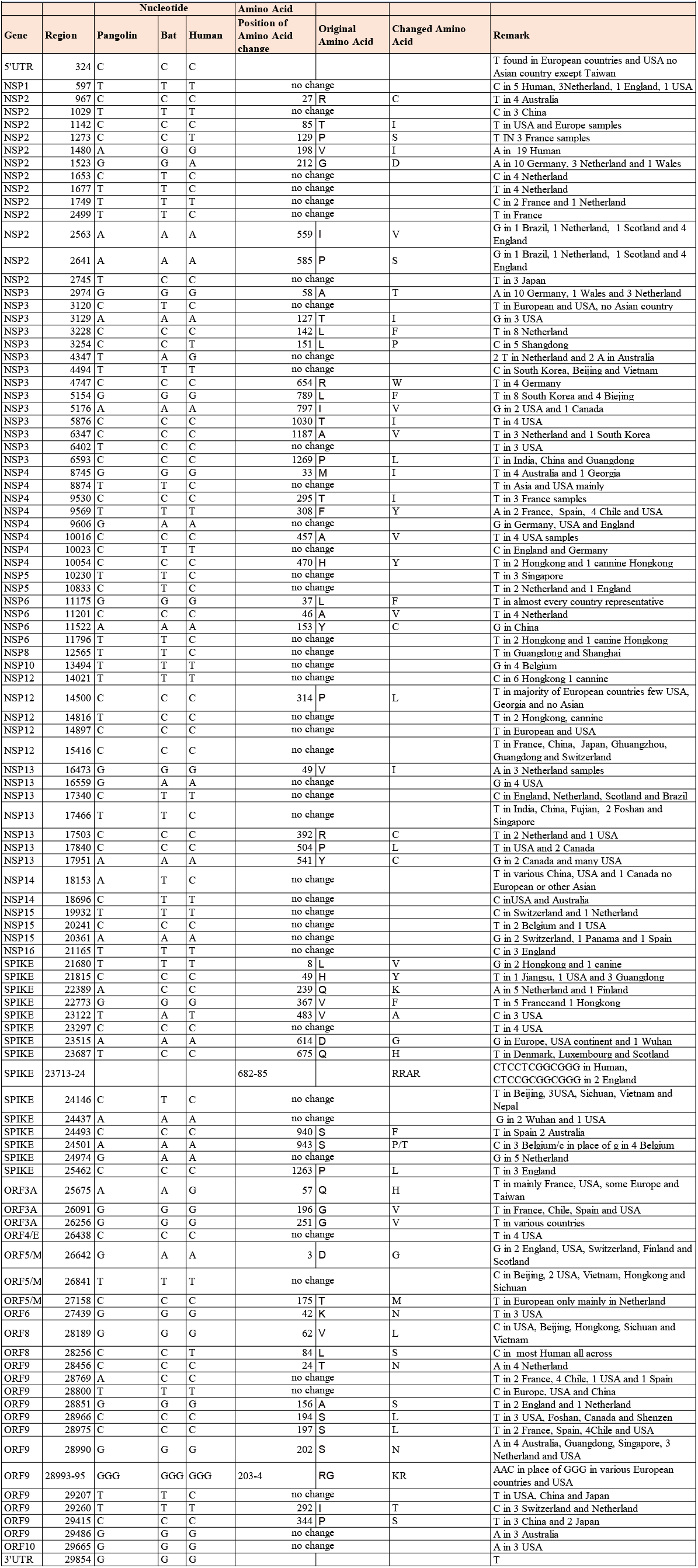
Distribution of synoymous, non-synonymous nucleotide changes and associated amino acid changes in different protein coding genes of hCoV genome

